# Differential Vulnerability of Stimulus-Locked and Persistent Gamma Oscillations: Implications in Schizophrenia

**DOI:** 10.64898/2026.05.29.728634

**Authors:** Daniel W. Chung, G. Bard Ermentrout

## Abstract

Working memory depends on gamma oscillations generated across sensory and prefrontal cortices. In sensory cortices such as primary visual cortex (V1), stimulus-locked gamma oscillations encode stimulus information, while in prefrontal cortex (PFC), persistent gamma oscillations maintain this information after the stimulus is removed. In schizophrenia (SZ), gamma power is reduced in both V1 and PFC, consistent with deficits in sensory encoding and working memory maintenance in the illness. These two regimes of gamma oscillations arise from a canonical microcircuit involving pyramidal neurons (PNs) and parvalbumin-expressing interneurons (PVIs). Yet, whether stimulus-locked and persistent gamma oscillations are similarly or differentially vulnerable to synaptic alterations within this circuit in SZ remains unknown. To investigate this question, we used a mean-field model of the PN-PVI circuit generating either stimulus-locked or persistent gamma oscillations. We then assessed the effects of three synaptic alterations found in SZ: lower excitatory drive to PVIs (E→I), lower inhibitory drive to PNs (I→E), and greater variability in E→I synaptic strength. Each alteration produced larger gamma power deficits in the persistent regime than in the stimulus-locked regime. When applied together, these alterations interacted synergistically to reduce gamma power in both regimes, with the persistent regime exhibiting a more pronounced deficit. Among the three parameters, E→I synaptic strength was the strongest contributor to the synergistic loss of gamma power. Two-dimensional bifurcation analyses further revealed that this differential vulnerability arises from a narrower margin of oscillatory stability in the persistent regime, where the parameter values producing maximum gamma power sit closer to the Hopf bifurcation boundary. Together, these findings identify the persistent regime as intrinsically more fragile than the stimulus-locked regime, with the implications for understanding regional patterns of synaptic pathology and cortical gamma oscillations with distinct dynamics in SZ.

**Author summary:** Working memory depends on stimulus-locked gamma oscillations in sensory cortices such as primary visual cortex (V1) for encoding stimulus information, and persistent gamma oscillations in prefrontal cortex (PFC) for maintaining this information after stimulus offset. In schizophrenia (SZ), gamma power is reduced in both V1 and PFC, and postmortem human brain studies suggest that the underlying synaptic alterations are more severe in V1 than in PFC. Our computational modeling results suggest that this regional pattern arises because persistent gamma oscillations are intrinsically more fragile than stimulus-locked gamma oscillations, so that smaller synaptic alterations are sufficient to disrupt gamma oscillations in PFC while larger alterations are required to produce comparable disruption in V1. Together, these findings give rise to a differential vulnerability model of cortical gamma oscillations in SZ, linking the regional patterns of synaptic pathology to the deficits in gamma oscillations observed across sensory and prefrontal cortices in the illness.

## Introduction

Working memory is a core cognitive process that relies on accurate encoding of sensory stimuli and maintenance of encoded information in the absence of stimuli [1–4]. These different components of working memory are thought to depend on neural oscillations at gamma frequency range (30 to 80 Hz) with distinct dynamics generated across different cortical areas [5]. For example, in sensory cortices such as primary visual cortex (V1), gamma oscillations are transiently evoked during stimulus presentation and terminate after stimulus offset (stimulus-locked gamma; [6–8]). In contrast, gamma oscillations in prefrontal cortex (PFC) persist after stimulus offset to maintain encoded information (persistent gamma; [9, 10]). Thus, understanding the neural substrates of working memory requires characterizing the circuit mechanisms that regulate stimulus-locked and persistent gamma oscillations.

Cortical gamma oscillations across sensory and prefrontal areas are thought to emerge from a canonical microcircuit involving excitatory pyramidal neurons (PNs) and inhibitory parvalbumin-expressing interneurons (PVIs) [11]. In this circuit, recurrently connected PNs provide excitatory drive to PVIs (E*→*I), which in turn provide phasic inhibition to PNs (I*→*E) that synchronizes PN firing at gamma frequency [12, 13]. Prior *in vivo* and *ex vivo* studies have established that the strengths of E*→*I and I*→*E synapses are the key determinants of cortical gamma power [14–19]. However, these studies did not directly compare stimulus-locked and persistent gamma regimes, and whether the two regimes are similarly or differentially regulated by E→I and I→E synaptic strength remains unexplored.

Studying the effects of E*→*I and I*→*E synaptic strengths on stimulus-locked and persistent gamma oscillations may provide insight into circuit-level mechanisms linking synaptic pathology to working memory dysfunction in schizophrenia (SZ). Prior postmortem studies in SZ have identified lower levels of molecular and/or structural markers of E*→*I [20–23] and I*→*E [24–28] synaptic strength across PFC and V1. Additionally, variability in the structural marker of E*→*I synaptic strength across individual PVIs was shown to be increased in the PFC of SZ [29], though this measure has not yet been assessed in V1. In line with these synaptic alterations, individuals with SZ exhibit reduced gamma power in visual cortex during stimulus encoding [30–32] and in PFC during the memory delay [33–35], and these deficits parallel impairments in early-stage sensory processing [36–39] and working memory performance [40–43] in the illness. Thus, determining whether E*→*I and I*→*E synaptic alterations differentially impact stimulus-locked and persistent gamma oscillations may clarify how a common synaptic pathology contributes to impairments in sensory encoding and working memory maintenance in SZ.

Computational modeling studies have provided key insights into how SZ-associated synaptic pathology affects gamma oscillation dynamics. For example, prior computational studies using networks of quadratic integrate-and-fire (QIF) neurons showed that lower levels of E*→*I and I*→*E synaptic strength can each disrupt gamma oscillations [28, 44–49]. In our previous work, we extended these findings by showing that greater variability in E*→*I synaptic strength also reduces gamma power, and demonstrated that these three synaptic alterations interact synergistically, with their combined effect on gamma power exceeding the sum of their individual effects [29]. Building on a growing body of literature using mean-field reductions of QIF networks [50–55], we further validated these individual and synergistic effects at the population level using a mean-field model designed to represent the collective dynamics of PNs and PVIs [56]. Together, these studies establish a computational framework for examining how synaptic alterations affect cortical gamma oscillation dynamics in SZ, motivating us to apply this approach to the distinct gamma oscillations supporting sensory encoding and memory maintenance.

In this study, we utilized a computational modeling approach to investigate whether and how stimulus-locked and persistent gamma oscillations are differentially affected by synaptic alterations implicated in SZ. Using a mean-field model of the PN-PVI circuit capable of generating either gamma oscillation regime, we systematically varied three SZ-associated synaptic parameters: E*→*I strength, I*→*E strength, and variability in E*→*I strength. We then assessed how each parameter individually and in combination affected gamma power in each regime, and identified which parameter contributed most strongly to the combined effect. Finally, we performed a series of two-dimensional bifurcation analyses to identify the dynamical mechanisms underlying the differential effects observed between the two regimes. Together, these analyses introduce a differential vulnerability model in which persistent gamma oscillations are intrinsically more fragile than stimulus-locked gamma oscillations, accounting for how a common set of synaptic alterations of differing magnitude gives rise to gamma oscillation deficits with distinct dynamics across sensory and prefrontal cortices in SZ.

## Materials and methods

### Mean-Field Model Description

In this study, we utilized a mean-field model of PN and PVI populations, which was derived from a network of quadratic integrate-and-fire (QIF) neurons in our recent work [56]. The QIF network consists of all-to-all connected regular-spiking PNs and fast-spiking PVIs coupled via AMPA, NMDA, and GABA synapses, forming a canonical pyramidal–interneuron gamma (PING) architecture that supports gamma oscillations through reciprocal excitatory–inhibitory interactions (Fig. 1A). Using the Ott–Antonsen ansatz [50, 57], the QIF network is reduced to a set of exact mean-field equations describing the mean firing rates and membrane potentials of the PN and PVI populations as follows:

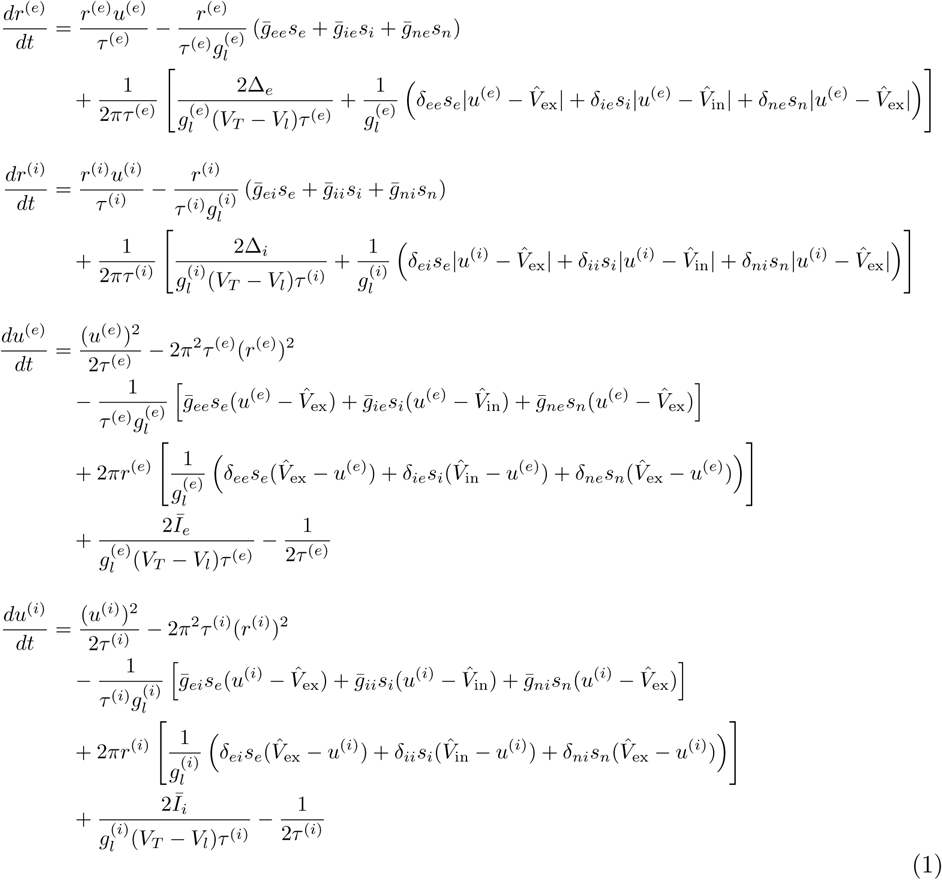

**Fig 1.**
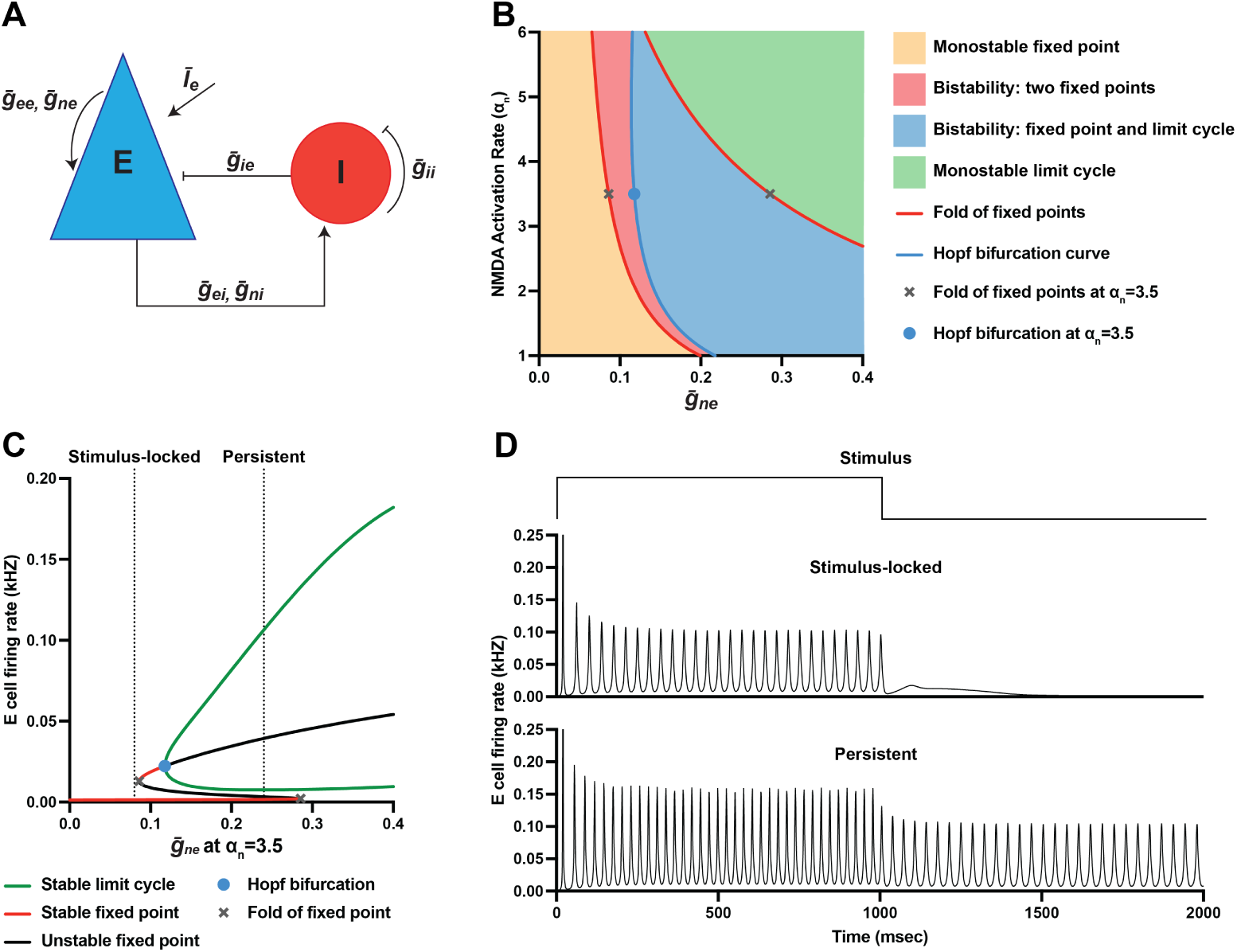
Emergence of distinct gamma oscillation regimes in a mean-field model of PING network. A. Schematic of the model network architecture consisting of reciprocally connected PNs and PVIs, with AMPA, NMDA, and GABA synapses. B. Two-dimensional bifurcation diagram showing the dynamical regimes of the mean-field model as a function of mean NMDA conductance among PNs (*ḡ_ne_*, x-axis) and NMDA activation rate constant(*a_n_*, y-axis). Regimes are separated by a fold of fixed points (red) and Hopf bifurcation curve (blue), revealing four qualitatively distinct regions: monostable fixed point (orange), bistability of two fixed points (red), bistability of a fixed point and a limit cycle (blue), and monostable limit cycle (green). C. One-dimensional bifurcation diagram showing network dynamics as a function of *ḡ_ne_* at fixed *a_n_* = 3.5. A fold and Hopf bifurcation points that correspond to those in panel B are marked by an “X” and blue circle, respectively. Vertical dotted lines indicate *ḡ_ne_* = 0.08 (left), which falls within the monostable fixed point regime associated with stimulus-locked gamma oscillations, and *ḡ_ne_* = 0.24 (right), which lies within the bistable regime of a fixed point and a limit cycle that supports persistent gamma oscillations. D. Simulated network activity demonstrating stimulus-locked gamma oscillations at *ḡ_ne_* = 0.08 (top) and persistent gamma oscillations at *ḡ_ne_* = 0.24 (bottom), both at *a_n_* = 3.5. External current was applied to PNs during the first 1000 ms of each 2000 ms simulation.

In these equations, superscripts (*e*) and (*i*) denote parameters for the PN and PVI populations, respectively. The variables *r* and *u* represent the mean firing rates and mean membrane potentials, respectively, of each population. The leak conductance is denoted by *g_l_*, and the membrane time constant *τ* is defined as the membrane capacitance divided by the leak conductance. The voltage threshold and the leak reversal potential are denoted by *V_T_* and *V_l_*, respectively. The reversal potentials for excitatory and inhibitory synaptic currents are denoted by *V*^^^_ex_ and *V*^^^_in_, respectively.

### Synaptic Connectivity

The external inputs to each population follow a Lorentzian distribution with mean of *Ī* and half-width of Δ. Synaptic connections are parameterized using *ḡ_xy_* and *δ_xy_*, which denote the mean and the half-width, respectively, of the Lorentzian distribution of synaptic conductance from presynaptic population *x* to postsynaptic population *y*. In this model, PNs are recurrently connected via AMPA- (*ḡ_ee_*) and NMDA-mediated (*ḡ_ne_*) synapses (Fig.1A). PNs also provide AMPA- (*ḡ_ei_*) and NMDA-mediated (*ḡ_ni_*) synaptic inputs to PVIs, which in turn provide GABAergic inhibition to PNs (*ḡ_ie_*) and to other PVIs (*ḡ_ii_*).

### Synaptic Gating Dynamics

The synaptic gating variables *s_e_*, *s_i_*, and *s_n_* represent the fraction of open AMPA, GABA, and NMDA receptors, respectively. These variables evolve according to:

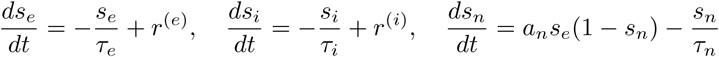

Here, *τ_e_*, *τ_i_*, and *τ_n_* are decay time constants for AMPA, GABA, and NMDA synapses, respectively, and *a_n_* is the NMDA activation rate constant.

### Model Validation

The mean-field model used in this study has been validated in our previous work by demonstrating close agreement with the underlying QIF network model of PNs and PVIs connected via AMPA-, NMDA-, and GABA-mediated synapses [56]. Both models exhibited overlapping gamma-frequency dynamics in response to external input, with closely matched firing patterns, gamma power, and peak frequency. Notably, although our model of the NMDA synapse omits the voltage-dependent Mg^2+^ block of NMDA receptors and treats *s_e_* and *s_n_* as statistically independent in deriving the mean-field equations, the close agreement with the underlying QIF network that also included NMDA synapses supports the validity of these approximations.

### Bifurcation and Numerical Simulation Procedures

We used XPPAUT to perform bifurcation analysis of the mean-field model. The full system of equations was implemented in XPPAUT, and AUTO was used to numerically continue fixed points and limit cycles as model parameters were varied.

For time-domain analysis, we numerically simulated the mean-field equations in MATLAB using an explicit Euler integration scheme with a fixed time step. Each simulation lasted 2000 ms, during which a step input current was applied to the PN population from 0 to 1000 ms to simulate transient external stimulation. The resulting firing rate of PNs, *r*^(^*^e^*^)^, was analyzed using MATLAB’s pwelch function to compute the power spectral density (PSD). To quantify stimulus-locked gamma power, PSD was computed over the stimulation period (0–1000 ms), while persistent gamma power was assessed by computing PSD during the post-stimulation period (1000–2000 ms). The values for all mean-field parameters are summarized in Table 1.

**Table 1.** List of parameters for the mean-field model.

| Parameter | Value |
| --- | --- |
| $\tau^{(e)}$ | 10 ms |
| $\tau^{(i)}$ | 5 ms |
| $g_l^{(e)}$ | 0.1 mS |
| $g_l^{(i)}$ | 0.2 mS |
| $V_l$ | −65 mV |
| $V_T$ | −50 mV |
| $\hat{V}_{\text{ex}}$ | 0 mV |
| $\hat{V}_{\text{in}}$ | −70 mV |
| $\tau_e$ | 2 ms |
| $\tau_i$ | 7 ms |
| $\tau_n$ | 80 ms |
| $a_n$ | Varied from 1 to 6 in Fig 1B; $3.5 \text{ ms}^{-1}$ in all other figures. |
| $\bar{g}_{ee}$ | 0.5 in all figures. |
| $\bar{g}_{ne}$ | Varied from 0 to 0.4 in Fig 1B, 1C; 0.08 for stimulus-locked regime, 0.24 for persistent regime in all other figures. |
| $\bar{g}_{ei}$ | Varied from 0 to 8 in Fig 2A–C; varied from 0 to 1 in Fig 4A, 4B; varied from healthy baseline ($\bar{g}_{ei,h} = 0.55$) to SZ state ($\bar{g}_{ei,sz} = 0.44$) in Figs 4C, 4D, 5, 6; 0.5 in all other figures. |
| $\bar{g}_{ni}$ | Varied from 0 to 0.3 in Fig 2D–F; varied from 0 to 0.1 in Fig 4A, 4B (linked to $\bar{g}_{ei}$ at a 1:10 ratio); varied from healthy baseline ($\bar{g}_{ni,h} = 0.055$) to SZ state ($\bar{g}_{ni,sz} = 0.044$) in Figs 4C, 4D, 5, 6; 0.05 in all other figures. |
| $\bar{g}_{ie}$ | Varied from 0 to 3 in Fig 2G–I; varied from 0 to 1 in Fig 4A, 4B; varied from healthy baseline ($\bar{g}_{ie,h} = 0.5$ for stimulus-locked, 0.45 for persistent) to SZ state ($\bar{g}_{ie,sz} = 0.4$ for stimulus-locked, 0.36 for persistent) in Figs 4C, 4D, 5, 6; 0.5 in all other figures. |
| $\bar{g}_{ii}$ | 0.5 in all figures. |
| $\delta_{ee}$ | 0.02 in all figures. |
| $\delta_{ne}$ | 0.02 in all figures. |
| $\delta_{ei}$ | Varied from 0 to 0.3 in Fig 3A–C; varied from healthy baseline ($\delta_{ei,h} = 0.02$) to SZ state ($\delta_{ei,sz} = 0.024$) in Figs 4C, 4D, 5, 6; 0.02 in all other figures. |
| $\delta_{ni}$ | Varied from 0 to 0.3 in Fig 3D–F; varied from healthy baseline ($\delta_{ni,h} = 0.02$) to SZ state ($\delta_{ni,sz} = 0.024$) in Figs 4C, 4D, 5, 6; 0.02 in all other figures. |
| $\delta_{ie}$ | 0.02 in all figures. |
| $\delta_{ii}$ | 0.02 in all figures. |
| $\bar{I}_e$ | 1.4 during stimulation; 0 otherwise. |
| $\bar{I}_i$ | 0 in all figures. |
| $\Delta_e$ | 0.05 in all figures. |
| $\Delta_i$ | 0.2 in all figures. |

## Results

### Bifurcation Analysis of Gamma Oscillation Regimes

Prior studies have established that the slow kinetics of NMDA conductance in recurrent excitatory synapses among PNs is critical for generating sustained neural activity after stimulus offset [58, 59]. Guided by this evidence, we performed a two-dimensional bifurcation analysis varying the mean NMDA conductance among PN population (*ḡ_ne_*) and the NMDA activation rate (*a_n_*) to identify the conditions under which the network supports stimulus-locked versus persistent gamma oscillations (Fig.1B).

This analysis revealed that as *ḡ_ne_* increases at fixed *a_n_*, the system transitions through four qualitatively distinct dynamical regimes. At low *ḡ_ne_* values, the network exhibited a monostable fixed-point regime (Fig.1B orange area). In this regime, the network was at an asynchronous low-activity state and produced gamma oscillations only during the presence of external stimulation, consistent with how stimulus-locked gamma oscillations emerge only in response to external input. As *ḡ_ne_* increased, the system entered a bistable regime with two fixed points (Fig.1B red area), in which the network could be pushed into a sustained, asynchronous high-activity state even after stimulus offset. With further increases in *ḡ_ne_*, the system transitioned into a bistable regime composed of a stable fixed point and a stable limit cycle (Fig.1B blue area). In this regime, gamma oscillations could be sustained after the removal of external stimulation, consistent with the dynamical property of persistent gamma oscillations. At high values of *ḡ_ne_*, the network entered a monostable limit cycle regime (Fig.1B green area) in which gamma oscillations were generated independent of external stimulation.

The boundaries between these regimes were defined by two types of bifurcation curves. First, the monostable fixed-point regime was bounded on the right by a fold of fixed points. The adjacent bistable regime with two fixed points emerged between this fold and a Hopf bifurcation curve. Furthermore, the bistable regime with a fixed point and a limit cycle was bounded on the left by the Hopf bifurcation curve and on the right by a second fold of fixed points. Finally, the monostable limit cycle regime lay beyond this second fold. These bifurcation-defined transitions tracked the system’s progression as *ḡ_ne_* increased, moving from stimulus-locked to persistent and ultimately to self-sustained gamma oscillations.

### Biologically Constrained Parameter Selection

To select biologically informed values of *ḡ_ne_* for simulating stimulus-locked and persistent gamma oscillations, we drew on findings linking the GluN2B subunit of the NMDA receptor to regional differences in recurrent excitatory synaptic function. In rodents, GluN2B is enriched in PNs relative to PVIs [60], and its slow deactivation kinetics are thought to be crucial for enabling the temporal summation necessary for sustained neural activity [61]. Consistent with this role, GluN2B mRNA levels in humans are approximately threefold higher in PFC than V1 [62], providing a biologically grounded basis for setting the *ḡ_ne_* value for the stimulus-locked and persistent regimes.

Accordingly, we searched the bifurcation diagram for values of *ḡ_ne_* that differed by approximately a factor of three and corresponded to dynamical regimes supporting either stimulus-locked or persistent gamma oscillations. At a fixed NMDA activation rate of *a_n_* = 3.5, *ḡ_ne_* = 0.08 placed the network in a monostable fixed-point regime that produced stimulus-locked gamma oscillations, while *ḡ_ne_* = 0.24 produced persistent gamma oscillations through bistability between a fixed point and a limit cycle (Fig. 1C,D). These values were used to simulate stimulus-locked and persistent gamma oscillations in all subsequent analyses.

### Differential Effect of E*→*I and I*→*E Synaptic Strength on Stimulus-Locked and Persistent Gamma Oscillations

Multiple in vivo studies have shown that excitatory drive to PVIs is a key determinant of cortical gamma power [15, 17–19]. In our previous modeling work, we found that increasing AMPA-mediated E*→*I conductance (*ḡ_ei_*) has an inverted U-shaped relationship with gamma power, whereas increasing NMDA-mediated E*→*I conductance (*ḡ_ni_*) leads to a monotonic reduction in gamma power [29, 56]. However, whether these effects differ between stimulus-locked and persistent gamma oscillations remains unclear.

To address this question, we first examined how changes in *ḡ_ei_* affect gamma power in the stimulus-locked and persistent regimes (Fig. 2A-C). In both regimes, gamma power increased sharply as *ḡ_ei_* rose from zero, peaking at *ḡ_ei_* = 0.7 for stimulus-locked regime and *ḡ_ei_* = 0.6 for persistent regime (Fig. 2A). Further increases in *ḡ_ei_* led to a progressive decline in gamma power, with a steeper drop in the persistent regime, reaching near-zero by *ḡ_ei_* = 2, compared to *ḡ_ei_* = 4 in the stimulus-locked regime. Consistent with these trends, bifurcation analysis revealed a broader oscillatory window for the stimulus-locked regime, with a Hopf bifurcation at *ḡ_ei_* = 0.37 indicating the onset of oscillations and a second Hopf bifurcation at *ḡ_ei_* = 7.17 marking their offset (Fig. 2B). In contrast, the persistent regime exhibited the same onset at *ḡ_ei_* = 0.37 but a much earlier offset at *ḡ_ei_* = 1.41 (Fig. 2C). These results indicate that the dynamic range of *ḡ_ei_* supporting gamma oscillations is markedly broader in the stimulus-locked regime compared to the persistent regime.

**Fig 2.**
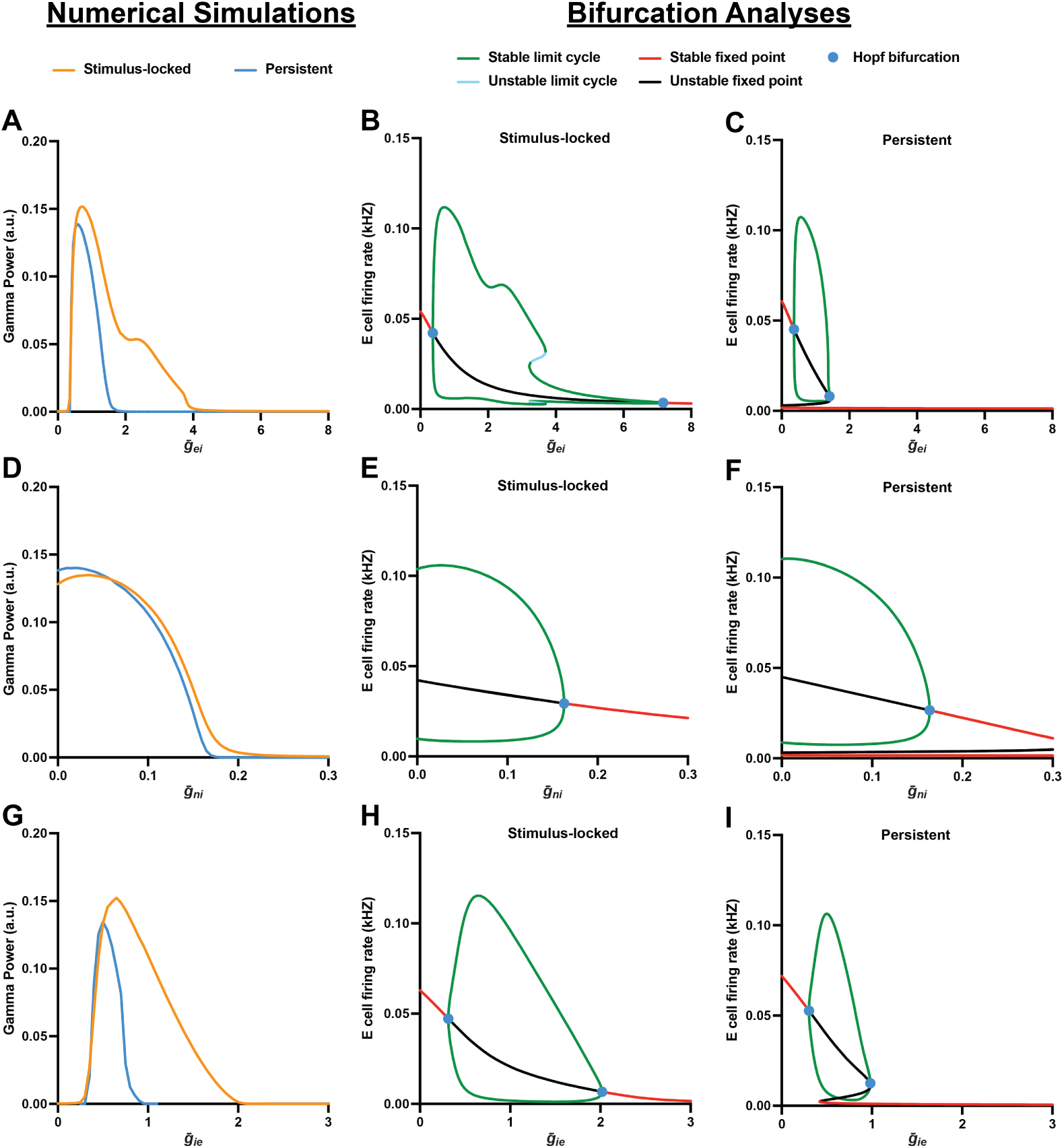
Differential effects of E→I and I→E synaptic strength on stimulus-locked and persistent gamma oscillations. A, D, G. Numerical simulations showing gamma power as a function of *ḡ_ei_* (A), *ḡ_ni_* (D), and *ḡ_ie_* (G) for the stimulus-locked (orange) and persistent (blue) regimes. B, E, H. One-dimensional bifurcation diagrams for the stimulus-locked regime as a function of *ḡ_ei_* (B), *ḡ_ni_* (E), and *ḡ_ie_* (H). C, F, I. One-dimensional bifurcation diagrams for the persistent regime as a function of *ḡ_ei_* (C), *ḡ_ni_* (F), and *ḡ_ie_* (I). In the bifurcation diagrams, stable and unstable limit cycles are denoted by green and light blue curves, respectively, stable and unstable fixed points by red and black curves, respectively, and Hopf bifurcation points by blue circles. For *ḡ_ei_* and *ḡ_ie_*, the oscillatory window of the persistent regime is narrower than that of the stimulus-locked regime, whereas for *ḡ_ni_*, the oscillatory windows are comparable across both regimes. Notably, in the persistent regime, the bifurcation structure of *ḡ_ie_* (I) differs qualitatively from that of *ḡ_ei_* (C): bistability between a stable fixed point and a limit cycle is present throughout the oscillatory window for *ḡ_ei_*, but emerges only above a fold of fixed points at 0.41 for *ḡ_ie_*.

We next examined how changes in *ḡ_ni_* influence gamma oscillations in the stimulus-locked and persistent regimes (Fig. 2D-F). In both regimes, gamma power progressively declined as *ḡ_ni_* increased from zero, reaching near-zero levels at *ḡ_ni_* = 0.22 in the stimulus-locked regime and *ḡ_ni_* = 0.18 in the persistent regime (Fig. 2D). Consistent with these trends, bifurcation analysis revealed similar limits of oscillatory stability, with Hopf bifurcations marking the offset of oscillations at *ḡ_ni_* = 0.16 in the stimulus-locked regime (Fig. 2E) and *ḡ_ni_* = 0.14 in the persistent regime (Fig. 2F). These findings suggest that the impact of *ḡ_ni_* on gamma oscillations is comparable across stimulus-locked and persistent regimes.

In addition to excitatory drive to PVIs, inhibitory drive from PVIs to pyramidal neurons has also been implicated as a key regulator of cortical gamma power [63, 64]. In our previous modeling work, we showed that increasing I*→*E synaptic strength (*ḡ_ie_*) leads to an inverted U-shaped relationship with gamma power [29, 56], similar to that observed for *ḡ_ei_*. To determine whether this relationship differs between stimulus-locked and persistent gamma oscillations, we examined the effects of varying *ḡ_ie_* in both regimes.

In both regimes, gamma power sharply increased as *ḡ_ie_* rose from zero, peaking at *ḡ_ie_* = 0.65 in the stimulus-locked regime and *ḡ_ie_* = 0.5 in the persistent regime (Fig. 2G). Beyond these peaks, gamma power progressively declined in both regimes, with a steeper drop in the persistent regime that reached near-zero levels by *ḡ_ie_* = 0.9 compared to *ḡ_ie_* = 2 in the stimulus-locked regime. Consistent with these trends, bifurcation analysis revealed a broader oscillatory window for the stimulus-locked regime, with a Hopf bifurcation at *ḡ_ie_* = 0.31 indicating the onset of oscillations and a second Hopf bifurcation at *ḡ_ie_* = 2.02 marking their offset (Fig. 2H). In contrast, the persistent regime exhibited a slightly later onset at *ḡ_ie_* = 0.36 and an earlier offset at *ḡ_ie_* = 1.41 (Fig. 2I). Notably, within this oscillatory window, bistability between a fixed point and a limit cycle emerged only above a fold at *ḡ_ie_* = 0.41, below which the network lacked a stable fixed point and oscillated continuously. Together, these findings indicate that the dynamic range of *ḡ_ie_* supporting gamma oscillations is narrower in the persistent regime than in the stimulus-locked regime.

### Comparable Effect of Synaptic Variability on Stimulus-Locked and Persistent Gamma Oscillations

We previously showed that PVIs receive excitatory input with variable synaptic strength, and that increasing this variability monotonically reduces gamma power [29, 56]. To determine whether this effect differs between stimulus-locked and persistent gamma oscillations, we assessed how gamma power changes in response to increasing the half-width of AMPA-mediated (*δ_ei_*) and NMDA-mediated (*δ_ni_*) E*→*I conductance across the PVI population.

Increasing *δ_ei_* progressively reduced gamma power in both the stimulus-locked and persistent regimes, with both reaching near-zero levels at *δ_ei_* = 0.2 (Fig. 3A). Consistent with this trend, bifurcation analysis revealed Hopf bifurcation points marking the offset of oscillations at *δ_ei_* = 0.19 in the stimulus-locked regime (Fig. 3B) and *δ_ei_* = 0.24 in the persistent regime (Fig. 3C). Similarly, increasing *δ_ni_* led to a progressive reduction in gamma power in both regimes, reaching near-zero levels near *δ_ni_* = 0.05 (Fig. 3D). Bifurcation analysis further showed Hopf bifurcation points marking the offset of oscillations at *δ_ni_* = 0.05 in both regimes (Fig. 3E,F). These results suggest that both stimulus-locked and persistent gamma oscillations are comparably sensitive to variability in excitatory synaptic input across PVIs.

**Fig 3.**
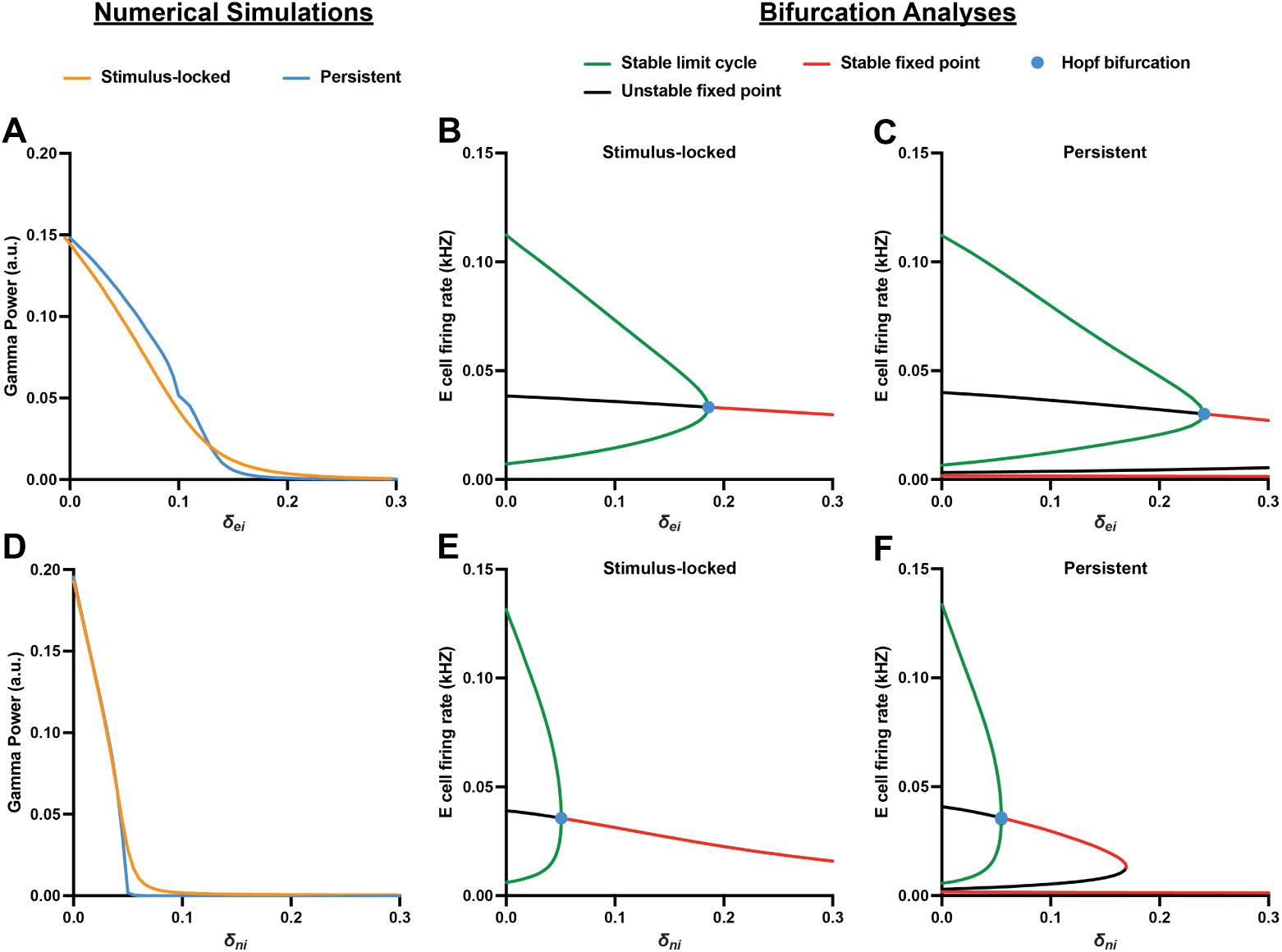
Comparable effects of E→I synaptic variability on stimulus-locked and persistent gamma oscillations. A, D. Numerical simulations showing gamma power as a function of *δ_ei_* (A) and *δ_ni_* (D) for the stimulus-locked (orange) and persistent (blue) regimes. B, E. One-dimensional bifurcation diagrams for the stimulus-locked regime as a function of *δ_ei_* (B) and *δ_ni_* (E). C, F. One-dimensional bifurcation diagrams for the persistent regime as a function of *δ_ei_* (C) and *δ_ni_* (F). In the bifurcation diagrams, stable limit cycles are denoted by green curves, stable and unstable fixed points by red and black curves, respectively, and Hopf bifurcation points by blue circles. For both *δ_ei_* and *δ_ni_*, increasing synaptic variability monotonically reduces gamma power, and the oscillatory windows are comparable across the stimulus-locked and persistent regimes.

### Differential Effect of SZ-Associated Synaptic Alterations on Stimulus-Locked and Persistent Gamma Oscillations

Prior postmortem studies in SZ have identified alterations in molecular and structural markers consistent with reduced E*→*I [20–23] and I*→*E [24–28] synaptic strength within PN-PVI microcircuits in PFC and V1, and increased variability in E*→*I strength across PVIs in the PFC [29]. In our prior modeling work, these alterations interacted synergistically to reduce gamma power [29, 56], motivating us to test whether the same set of perturbations produces similar or distinct effects on stimulus-locked versus persistent gamma oscillations.

First, we sought to identify a set of parameter values to represent the healthy state in each regime. To establish this parameter set, we simulated the effect of interaction between *ḡ_ei_*, *ḡ_ni_*, and *ḡ_ie_* on gamma power and searched for parameter values that produced maximal gamma power (Fig. 4A,B). In these simulations, we modeled changes in the total E*→*I synaptic strength by adjusting *ḡ_ei_* and *ḡ_ni_* simultaneously while maintaining a fixed ratio of 1:10 to reflect the NMDA-to-AMPA ratio empirically observed in PVIs [46]. Additionally, we set *δ_ei_* and *δ_ni_* at 0.02, corresponding to the lower end of the range for these parameters that results in a linear decrease in gamma power. Because the underlying network dynamics differ between the two regimes, we performed this optimization independently for both the stimulus-locked and persistent regimes. The parameter values that produced maximum gamma power, representing the healthy baseline (*h*) state, occurred for each regime at:

**Fig 4.**
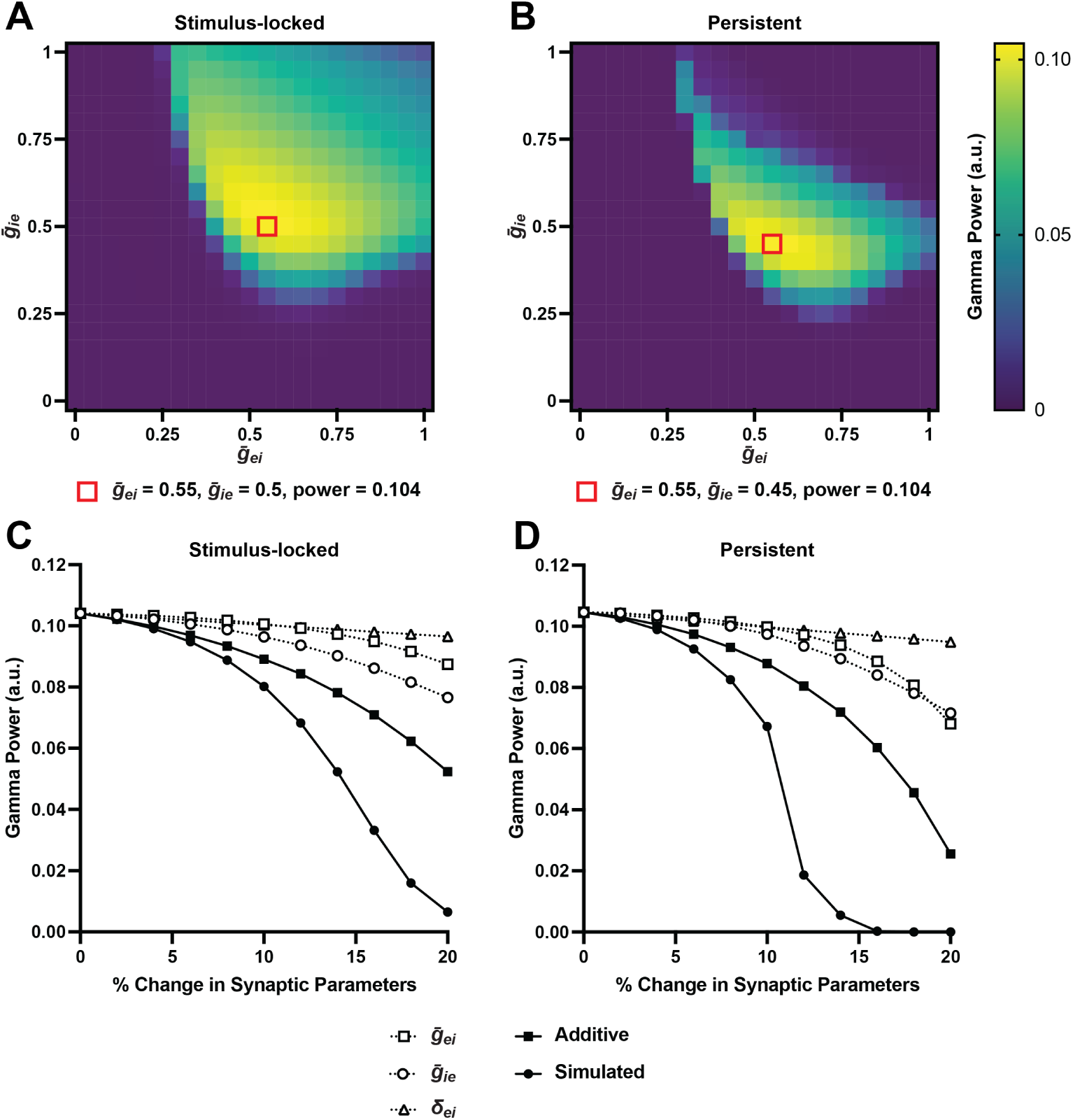
Differential effects of SZ-associated synaptic alterations on stimulus-locked and persistent gamma oscillations. A, B. Heatmaps showing gamma power as a function of *ḡ_ei_* (x-axis) and *ḡ_ie_* (y-axis) for the stimulus-locked (A) and persistent (B) regimes. Red squares indicate the parameter sets that produced maximum gamma power, representing the healthy baseline for each regime (*ḡ_ei_* = 0.55, *ḡ_ie_* = 0.5 for stimulus-locked; *ḡ_ei_* = 0.55, *ḡ_ie_* = 0.45 for persistent). Both regimes yielded an identical maximum gamma power of 0.104. C, D. Gamma power as a function of progressive SZ-associated synaptic alterations (0–20%) for the stimulus-locked (C) and persistent (D) regimes. Dotted lines show individual effects of reducing *ḡ_ei_* (squares), reducing *ḡ_ie_* (circles), and increasing *δ_ei_* (triangles). Solid line with filled squares shows the expected additive sum of individual effects, and solid line with filled circles shows the simulated combined effect. In both regimes, the simulated combined effect exceeds the expected additive sum, indicating synergistic interaction. In the stimulus-locked regime, a full 20% alteration resulted in a 94% reduction of gamma power, doubling the expected additive deficit of 50%. In contrast, the persistent regime reached this same 94% reduction when synaptic parameters were altered by only 14%, more than tripling the expected additive deficit of 31% at that point. In all panels, *ḡ_ei_* and *δ_ei_* were varied jointly with their NMDA counterparts *ḡ_ni_* and *δ_ni_*, respectively.

#### The stimulus-locked regime

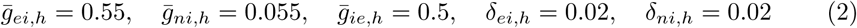

#### The persistent regime

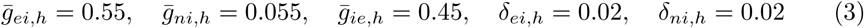

Notably, these parameter sets yielded an identical maximum gamma power of 0.104 for both the stimulus-locked and persistent regimes, ensuring that the subsequent simulation results are due to the intrinsic dynamics of each regime rather than differences in their initial gamma power.

We then defined a parameter set representing the SZ state for each regime based on the studies that quantified specific molecular or structural markers most directly mapping onto each of the three modeled synaptic alterations (Table 2). For E*→*I synaptic strength, we drew on two complementary markers. The density of VGlut1^+^/PSD95^+^ puncta on PVIs provides a direct morphological measure of E*→*I strength and is lower in PFC of SZ [21], but this measure has not yet been quantified in V1. As a complementary marker, the shift in ErbB4 splicing from major to minor variants has been shown to reduce the number of excitatory inputs to PVIs [65] and to correlate with VGlut1^+^/PSD95^+^ puncta density onto PVIs in PFC of SZ [21]. This abnormal splicing shift was evident in both PFC and V1 of SZ in a recent cross-regional study [23], supporting reductions in *ḡ_ei_* and *ḡ_ni_* in both stimulus-locked and persistent regimes. For I*→*E synaptic strength, we used a composite measure of PV inhibitory terminal strength, defined by VGAT^+^/PV^+^ puncta density combined with protein levels of the GABA-synthesizing enzymes GAD65 and GAD67 within these puncta. A cross-regional study found that this composite measure is lower in both PFC and V1 of SZ [26], supporting reduction in *ḡ_ie_* in both regimes. For the variability in E*→*I synaptic strength, we used the coefficient of variation in VGlut1^+^ and PSD95^+^ puncta protein levels across PVIs, which is greater in PFC of SZ [29]. Although this measure has not yet been assessed in V1, we increased *δ_ei_* and *δ_ni_* in both regimes so that all three synaptic alterations could be evaluated together.

**Table 2.** Postmortem findings in SZ across PFC and V1, with their expected synaptic changes and corresponding parameters in the mean-field model. Magnitudes indicate the direction and percent change relative to unaffected comparison subjects. For the markers assessed in both regions, alterations are larger in V1 than in PFC.

| Postmortem findings in SZ | Expected synaptic change | PFC magnitude | V1 magnitude | Model parameter |
| --- | --- | --- | --- | --- |
| Lower VGlut1<sup>+</sup>/PSD95<sup>+</sup> puncta density in PVIs [21] | Lower E→I | −18% | Not assessed | $\bar{g}_{ei}, \bar{g}_{ni}$ |
| ErbB4 splicing shift from major to minor variants [23] | Lower E→I | −15% | −52% | $\bar{g}_{ei}, \bar{g}_{ni}$ |
| Composite measure of PV inhibitory terminal strength [26] | Lower I→E | −22% | −42% | $\bar{g}_{ie}$ |
| Greater variability in VGlut1 and PSD95 puncta level across PVIs [29] | Greater E→I variability | +20–28% | Not assessed | $\delta_{ei}, \delta_{ni}$ |

For the markers that have been assessed in both PFC and V1, the magnitude of alterations in SZ was consistently greater in V1 (Table 2). For the parameter set representing the SZ state, we applied a uniform 20% perturbation to the healthy baseline values in both regimes, based on PFC data where the magnitude of alterations ranged from 15% to 28% in SZ relative to unaffected subjects. We selected this approach for three reasons. First, applying the same magnitude across both regimes enabled direct comparison of how stimulus-locked and persistent gamma oscillations respond to identical perturbations. Second, using PFC-derived magnitudes rather than the larger V1 deficits avoided the risk of eliminating gamma oscillations in both regimes, which would prevent us from determining whether the two regimes are differentially affected by synaptic alterations. Third, applying a uniform magnitude across all three parameters ensured that no single alteration dominated the combined effect. These SZ-associated parameter values (*sz*) were:

#### For the stimulus-locked regime

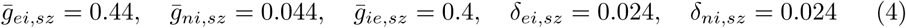

#### For the persistent regime

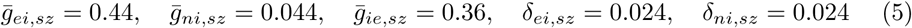

To systematically examine how the accumulation of these alterations affects gamma power in each regime, we simulated a linear transition from the healthy baseline of 0% alteration to the full SZ state representing the 20% alteration (Fig. 4C,D). For brevity, throughout subsequent analyses we refer only to the AMPA parameters *ḡ_ei_* and *δ_ei_*, with the understanding that their NMDA counterparts *ḡ_ni_* and *δ_ni_* are varied in parallel. We first examined how each synaptic alteration individually impacted gamma power. In the stimulus-locked regime, a 20% decrease in *ḡ_ie_* and *ḡ_ei_* reduced gamma power by 26% and 16%, respectively, while a 20% increase in *δ_ei_* reduced it by 7% (Fig. 4C). In the persistent regime, the same 20% alterations produced larger reductions, with *ḡ_ie_* and *ḡ_ei_* reducing gamma power by 31% and 35%, respectively, and *δ_ei_* reducing it by 9% (Fig. 4D).

We then investigated whether these synaptic alterations interact synergistically when applied in combination. In both regimes, the combined reduction in gamma power was greater than the sum of the individual effects, indicating synergistic interaction (Fig. 4C,D). In the stimulus-locked regime, simultaneously applying all three alterations at the full 20% level resulted in a 94% reduction of gamma power, nearly doubling the expected additive deficit of 50% (Fig. 4C). The persistent regime exhibited an even stronger synergy, reaching this same 94% reduction when the three parameters were altered by only 14%, at which point the deficit was more than triple the expected additive reduction of 31% (Fig. 4D). Together, these results indicate that the synergistic interaction among SZ-associated synaptic alterations is conserved across both regimes but is markedly stronger in the persistent regime than in the stimulus-locked regime.

To quantify the magnitude of synergistic interactions between these synaptic alterations, we calculated a synergy index, defined as the ratio of the simulated gamma power deficit to the expected additive deficit (Fig. 5A). An index value greater than 1 indicates a synergistic interaction, where the combined impact of synaptic changes is greater than the sum of their individual effects. In the stimulus-locked regime, the synergy index increased gradually as the synaptic parameters shifted away from the healthy baseline, reaching a peak of 2.2 at 16.5% alteration. At the full 20% alteration, the index slightly declined to 1.9, remaining above the additive threshold. In contrast, the persistent regime exhibited a markedly different synergy profile, with the synergy index increasing rapidly to a peak of 3.8 at only 12.5% alteration. Following this peak, the index declined sharply to 1.3 at the full 20% alteration.

**Fig 5.**
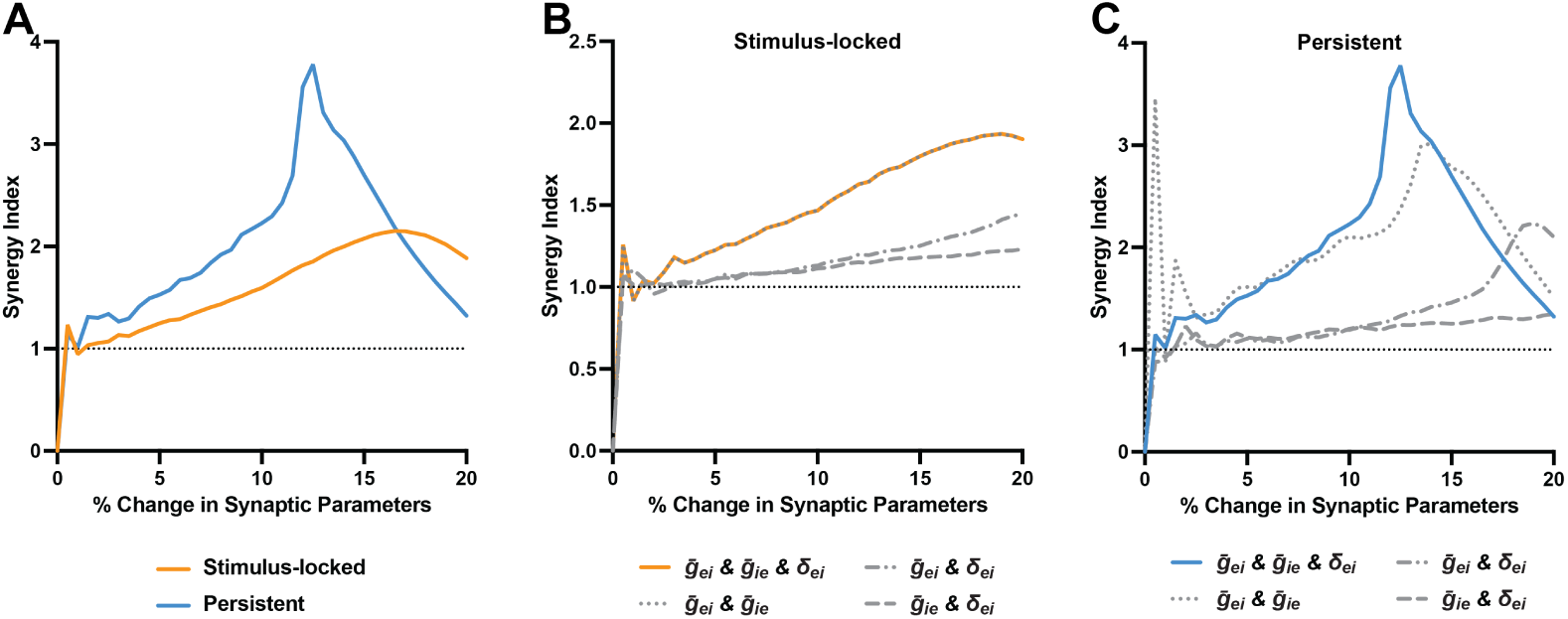
Synergy analysis of SZ-associated synaptic alterations in stimulus-locked and persistent gamma oscillations. A. Synergy index as a function of progressive SZ-associated synaptic alterations (0–20%) for the stimulus-locked (orange) and persistent (blue) regimes. The synergy index is defined as the ratio of the simulated gamma power deficit to the expected additive deficit, with values greater than 1 (dotted horizontal line) indicating synergistic interaction. The stimulus-locked regime reached a peak synergy index of 2.2 at 16.5% alteration, while the persistent regime reached a higher peak of 3.8 at only 12.5% alteration. B, C. Pairwise synergy analysis for the stimulus-locked (B) and persistent (C) regimes, showing the synergy index for the combined interaction of all three parameters (solid colored line) alongside the synergy indices for each pairwise combination: *ḡ_ei_* & *ḡ_ie_* (dotted), *ḡ_ei_* & *δ_ei_* (dash-dot), and *ḡ_ie_* & *δ_ei_* (dashed). In both regimes, *ḡ_ei_* is the strongest contributor to synergistic gamma power loss, followed by *ḡ_ie_*, with *δ_ei_* being the weakest. In the stimulus-locked regime, the synergy is driven almost entirely by the combined effect of *ḡ_ei_* and *ḡ_ie_*, with *δ_ei_* contributing minimally. In the persistent regime, the dominance of *ḡ_ei_* over *ḡ_ie_* is more pronounced, and *δ_ei_* plays a larger role than in the stimulus-locked regime. In all panels, *ḡ_ei_* and *δ_ei_* were varied jointly with their NMDA counterparts *ḡ_ni_* and *δ_ni_*, respectively.

To identify the primary drivers of the synergistic interactions observed in each regime, we assessed the synergy index while systematically isolating the interactions between only two of the three SZ-associated alterations (Fig. 5B,C). This allowed us to determine which synaptic parameter most strongly contributes to the overall synergistic reduction in gamma power. In the stimulus-locked regime, the synergy index for the combined interaction of all three parameters was almost entirely explained by the *ḡ_ie_* and *ḡ_ei_* pairing (Fig. 5B). The *ḡ_ei_* & *δ_ei_* and *ḡ_ie_* & *δ_ei_* pairings produced much lower synergy indices that remained relatively flat, suggesting that synergy is primarily driven by the combined reduction of both mean synaptic strengths. Between these two pairings, the *ḡ_ei_* & *δ_ei_* pairing produced a slightly higher synergy index than the *ḡ_ie_* & *δ_ei_* pairing, indicating that *ḡ_ei_* is a modestly stronger contributor than *ḡ_ie_*. In the persistent regime, the *ḡ_ei_* & *ḡ_ie_* pairing remained the strongest contributor but no longer fully accounted for the total synergistic effect (Fig. 5C), indicating that *δ_ei_* contributes more meaningfully to synergy in the persistent regime than in the stimulus-locked regime. Furthermore, the *ḡ_ei_* & *δ_ei_* pairing showed a more pronounced rise in synergy index compared to that in the stimulus-locked regime, while the *ḡ_ie_* & *δ_ei_* pairing remained similarly low across both regimes, indicating that the dominance of *ḡ_ei_* over *ḡ_ie_* is larger in the persistent regime. Together, these results indicate that *ḡ_ei_* is the strongest contributor to synergistic gamma power loss in both regimes, followed by *ḡ_ie_*, with *δ_ei_* being the weakest. In the persistent regime, however, the dominance of *ḡ_ei_* over *ḡ_ie_* is more pronounced and *δ_ei_* plays a larger role than in the stimulus-locked regime.

### Two-Dimensional Bifurcation Analyses of Regime-Specific Vulnerabilities

To understand the dynamical mechanisms underlying the differential effects of SZ-associated synaptic alterations on stimulus-locked and persistent gamma oscillations, we performed two-dimensional bifurcation analyses in three pairwise combinations of *ḡ_ei_*, *ḡ_ie_*, and *δ_ei_* (Fig. 6A-C). In each plane, two parameters were varied along the axes while the third parameter was held at either its healthy baseline (0% perturbation) or its full SZ state (20% perturbation). This produced two Hopf bifurcation curves that bound the oscillatory region: the healthy Hopf curve, traced with the third parameter at its healthy baseline, and the SZ Hopf curve, traced with the third parameter at its full SZ state. We then superimposed the SZ trajectory of the two axis parameters, progressing in 5% increments from their healthy baseline to their full SZ state. In the persistent regime, a fold of fixed points also appeared in each parameter plane, demarcating the boundary between the monostable limit cycle regime and the bistable regime supporting persistent gamma oscillations. The stimulus-locked regime, which lacks bistability, exhibited only Hopf bifurcation curves.

**Fig 6.**
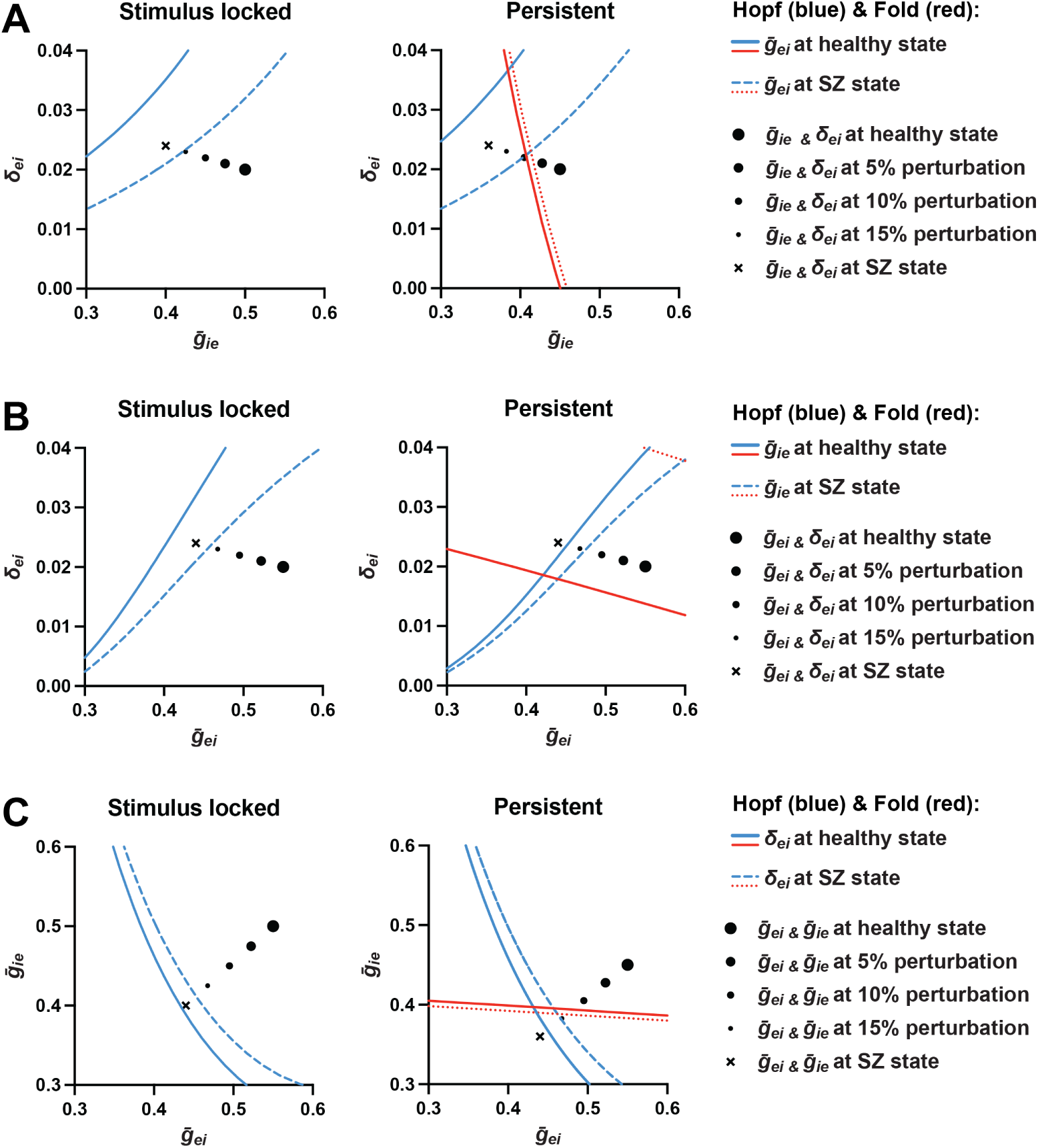
Two-dimensional bifurcation analyses of regime-specific vulnerabilities. A–C. Two-dimensional bifurcation analyses for the stimulus-locked (left) and persistent (right) regimes in three pairwise parameter planes: *ḡ_ie_* vs. *δ_ei_* (A), *ḡ_ei_* vs. *δ_ei_* (B), and *ḡ_ei_* vs. *ḡ_ie_* (C). Solid and dashed blue curves denote Hopf curves at the healthy and SZ-associated values of the third parameter, respectively. In the persistent regime, solid and dotted red curves denote folds of fixed points at the healthy and SZ-associated values of the third parameter, demarcating the boundary between the bistable persistent gamma regime and the monostable limit cycle regime. Black circles of decreasing size represent the SZ perturbation trajectory of the two axis parameters, progressing from the healthy baseline (largest circle) in 5% increments to the full SZ state (x). Across all three planes, the healthy baseline values of the two axis parameters sat closer to the Hopf curves and their SZ trajectory crossed the SZ Hopf curve at smaller perturbation magnitudes in the persistent regime than in the stimulus-locked regime. In the persistent regime, the trajectory also crossed the fold of fixed points prior to crossing the SZ Hopf curve in A and C, and in B the trajectory’s starting point already lay past the fold when *ḡ_ie_* was at its SZ state. In all panels, *ḡ_ei_* and *δ_ei_* were varied jointly with their NMDA counterparts *ḡ_ni_* and *δ_ni_*, respectively.

In the *ḡ_ie_* vs. *δ_ei_* plane, the third parameter *ḡ_ei_* was held at either its healthy or SZ state (Fig. 6A). The healthy baseline of the two axis parameters sat closer to both Hopf curves in the persistent regime than in the stimulus-locked regime. Consistent with this proximity, the SZ trajectory of the two axis parameters crossed the SZ Hopf curve at 10% perturbation in the persistent regime, compared to 15% in the stimulus-locked regime. In the persistent regime, the SZ trajectory also crossed the fold of fixed points before reaching the SZ Hopf curve. This earlier crossing, primarily driven by the decrease in *ḡ_ie_*, indicates that bistability was lost before the oscillation itself was eliminated.

In the *ḡ_ei_* vs. *δ_ei_* plane, the third parameter *ḡ_ie_* was held at either its healthy or SZ state (Fig. 6B). The healthy baseline of the two axis parameters sat closer to both Hopf curves in the persistent regime than in the stimulus-locked regime. Consistent with this proximity, the SZ trajectory of the two axis parameters crossed the SZ Hopf curve at 10–15% perturbation in the persistent regime, compared to 15–20% in the stimulus-locked regime. In the persistent regime, the fold of fixed points lay below the SZ trajectory when *ḡ_ie_* was at its healthy state, indicating that bistability was intact along the trajectory. When *ḡ_ie_* was at its SZ state, however, the trajectory’s starting point already lay in the monostable limit cycle regime, indicating that reduction of *ḡ_ie_* alone was sufficient to abolish bistability, consistent with Fig. 6A.

In the *ḡ_ei_* vs. *ḡ_ie_* plane, the third parameter *δ_ei_* was held at either its healthy or SZ state (Fig. 6C). The healthy baseline of the two axis parameters sat closer to both Hopf curves in the persistent regime than in the stimulus-locked regime. Consistent with this proximity, the SZ trajectory of the two axis parameters crossed the SZ Hopf curve at 15% perturbation in the persistent regime, compared to 15–20% in the stimulus-locked regime. As in Fig. 6A, the SZ trajectory in the persistent regime crossed the fold of fixed points before reaching the SZ Hopf curve.

Comparing the three parameter planes, the magnitude of Hopf-curve displacement by the third parameter from healthy to SZ states followed a consistent ordering, with *ḡ_ei_* producing the largest displacement, followed by *ḡ_ie_*, and *δ_ei_* producing the smallest, matching the synergy contribution hierarchy identified in the preceding analysis. Moreover, the difference in Hopf-curve displacement produced by *ḡ_ei_* versus *ḡ_ie_* was larger in the persistent regime than in the stimulus-locked regime, consistent with the larger gap in their synergy contributions. Together, these analyses reveal three dynamical mechanisms that distinguish the persistent regime from the stimulus-locked regime. First, the Hopf curves sit closer to the healthy baseline of the two axis parameters in the persistent regime, reflecting a narrower oscillatory window. As a result, the SZ trajectory crosses the SZ Hopf curve at smaller perturbation magnitudes, accounting for the earlier loss of gamma power. Second, the dominance of *ḡ_ei_* over *ḡ_ie_* in Hopf-shift magnitude is more pronounced in the persistent regime, accounting for the larger gap in their synergy contributions. Third, in the persistent regime, reduction of *ḡ_ie_* crosses the fold of fixed points before reaching the Hopf curve, abolishing bistability and transitioning the network from persistent gamma oscillations to continuous baseline oscillations.

## Discussion

In this study, we utilized a mean-field model of the PN-PVI circuit to investigate whether and how stimulus-locked and persistent gamma oscillations are differentially affected by synaptic alterations implicated in SZ. The model generated either stimulus-locked or persistent gamma oscillations based on empirical differences in NMDA receptor subunit expression between V1 and PFC. We then assessed the effects of three SZ-associated synaptic alterations: lower E*→*I synaptic strength (*ḡ_ei_* and *ḡ_ni_*), lower I*→*E synaptic strength (*ḡ_ie_*), and greater variability in E*→*I synaptic strength (*δ_ei_* and *δ_ni_*). Each alteration individually produced larger gamma power deficits in the persistent regime than in the stimulus-locked regime. When applied together, these alterations interacted synergistically to reduce gamma power in both regimes, with the persistent regime exhibiting an earlier and more pronounced collapse. In both regimes, the contribution to synergistic gamma power loss was greatest for lower E*→*I strength, intermediate for lower I*→*E strength, and smallest for greater E*→*I variability, though the dominance of lower E*→*I over lower I*→*E strength was more pronounced in the persistent regime. Two-dimensional bifurcation analyses revealed that the greater vulnerability of the persistent regime to synaptic alterations arises from a narrower intrinsic margin of oscillatory stability, while the parameter-specific contribution hierarchy reflects the magnitude of Hopf-curve displacement produced by each synaptic parameter. Additionally, in the persistent regime, reducing I*→*E synaptic strength eventually abolished bistability by crossing a fold of fixed points, transitioning the network from persistent to continuous baseline oscillations.

The central finding of this study is that persistent gamma oscillations are intrinsically more vulnerable than stimulus-locked gamma oscillations to SZ-associated synaptic alterations. This differential vulnerability was evident in both numerical simulations and bifurcation analyses. In the one-dimensional bifurcation analyses, the oscillatory window of the persistent regime was consistently narrower than that of the stimulus-locked regime along E*→*I and I*→*E strength. In the synergy analyses, this vulnerability manifested as an earlier and more pronounced collapse of gamma power in the persistent regime when SZ-associated alterations were applied in combination. The two-dimensional bifurcation analyses revealed the dynamical basis of this vulnerability: the healthy baseline values of the synaptic parameters that were chosen to produce the maximal gamma power sit closer to the Hopf bifurcation curve in the persistent regime than in the stimulus-locked regime. As a result, the SZ perturbation trajectory crosses the Hopf curve earlier in the persistent regime than in the stimulus-locked regime, so that smaller magnitudes of perturbations are sufficient to disrupt persistent gamma oscillations. This intrinsic difference in the margin of oscillatory stability provides the dynamical foundation for the differential vulnerability of stimulus-locked and persistent gamma oscillations to SZ-associated synaptic alterations.

These findings give rise to a differential vulnerability model of cortical gamma oscillations in SZ. Although gamma power is lower in both V1 and PFC in SZ, the present modeling results predict that the underlying synaptic pathology could differ in magnitude across these regions. Specifically, because the persistent regime is more easily disrupted than the stimulus-locked regime, relatively small synaptic alterations should be sufficient to disrupt persistent gamma oscillations in the PFC, whereas larger synaptic alterations would be required to disrupt stimulus-locked gamma oscillations in V1. Recent cross-regional postmortem studies provide empirical support for this prediction. First, a composite measure of PV basket cell inhibitory strength onto PNs is lower in V1 and PFC in SZ, with approximately twofold greater deficits in V1 than in PFC [26]. Second, the shift in ErbB4 splicing from major to minor variants, a marker of lower E*→*I synaptic strength onto PVIs [21], is larger in V1 than in PFC in SZ [23]. Consistent with these alterations, PV mRNA level, a marker of PVI activity [66], is lower in both regions in SZ with a larger deficit in V1 than in PFC [23, 27], reflecting a regional pattern of PVI activity that parallels the synaptic alterations. Together, these convergent patterns of empirical findings in SZ and the prediction of our computational model provide a framework for linking the region-specific magnitude of synaptic pathology to differential vulnerability of cortical gamma oscillations with distinct dynamics in SZ.

Our study also reveals that SZ-associated reduction of I*→*E strength can abolish the bistability required for persistent gamma oscillations. In the healthy persistent regime, bistability between a stable low-activity fixed point and a limit cycle allows the network to sit quietly before stimulation and then sustain gamma oscillations after stimulus offset. When I*→*E strength is reduced toward its SZ state, the network crosses a fold of fixed points, eliminating the stable low-activity fixed point and leaving only the limit cycle. In this state, the network oscillates continuously at baseline without requiring a stimulus to initiate the oscillation, representing a transition from task-related persistent gamma oscillations to resting-state gamma oscillations. Notably, elevated resting-state gamma power has been reported in the PFC of individuals with SZ [67, 68]. While multiple mechanisms could contribute to this elevation, the present modeling results raise the possibility that reduced I*→*E strength, by abolishing bistability and producing continuous baseline oscillations, could be one contributing factor.

Our analysis identified a modest role for greater variability in E*→*I strength in the synergistic disruption of gamma oscillations. Across individual perturbations, synaptic variability was consistently the weakest contributor of the three SZ-associated synaptic parameters examined. However, while the interaction between lower E*→*I and lower I*→*E strength fully accounted for the total synergy in the stimulus-locked regime, this interaction did not fully account for the total synergy in the persistent regime, indicating that synaptic variability plays a larger role in disrupting persistent gamma oscillations. These findings predict that increased synaptic variability should have a greater impact on gamma power in PFC than in V1. Consistent with this prediction, we previously reported increased coefficient of variation in excitatory synapse markers across individual PVIs in the PFC of SZ [29], though whether a similar increase is present in V1 remains to be examined.

Several limitations of this study should be acknowledged. First, we selected the NMDA conductance values that distinguish the stimulus-locked and persistent regimes based on the threefold difference in GluN2B mRNA expression between human V1 and PFC [62]. Because mRNA expression is an indirect measure of NMDA synaptic conductance, future work using more direct measures, such as NMDA receptor protein levels or electrophysiological recordings of NMDA currents, will help refine the NMDA conductance values appropriate for each regime. Second, our model represents PVIs as a single homogeneous population of PV basket cells that provide perisomatic inhibition to neighboring PNs. However, PVIs also include chandelier cells that instead target the axon initial segment of PNs [69], and PV basket cells themselves comprise distinct subpopulations with different connectivity profiles, target PN types, and plasticity rules [70]. Whether these subpopulations contribute differently to stimulus-locked and persistent gamma oscillations and if they are differentially affected in SZ will be important to investigate in future work. Third, our analysis focused on three synaptic parameters within the local PN-PVI circuit but several other mechanisms that shape cortical gamma oscillations are also disrupted in SZ. For example, thalamocortical connectivity [71, 72] and the activity of somatostatin-expressing interneurons [73] each contribute to generating cortical gamma oscillations and are thought to be disrupted in SZ [74–77]. Incorporating these elements into future models will be important for capturing the full complexity of cortical gamma oscillation dysfunction in the illness.

In conclusion, our computational modeling results identify persistent gamma oscillations as intrinsically more vulnerable than stimulus-locked gamma oscillations to SZ-associated synaptic alterations. This differential vulnerability model posits that the synaptic pathology underlying gamma oscillation dysfunction in SZ could vary in magnitude across cortical regions, with larger alterations required in sensory areas supporting the stimulus-locked regime than in prefrontal areas supporting the persistent regime, a prediction supported by multiple postmortem studies. Together, these results provide a cohesive account of how a shared synaptic disruption of differing magnitude across cortical regions can give rise to gamma oscillation deficits that diverge by region, oscillatory dynamics, and their contributions to working memory dysfunction in SZ.

